# MiSDEED: a synthetic multi-omics engine for microbiome power analysis and study design

**DOI:** 10.1101/2021.08.09.455682

**Authors:** Philippe Chlenski, Melody Hsu, Itsik Pe’er

## Abstract

**Summary:** MiSDEED is a command-line tool for generating synthetic longitudinal multi-omics data from simulated microbial environments. It generates relative-abundance timecourses under perturbations for an arbitrary number of samples and patients. All simulation parameters are exposed to the user to facilitate rapid power analysis and aid in study design. Users who want additional flexibility may also use MiSDEED as a Python package.

**Availability and implementation:** MiSDEED is written in Python and is freely available at https://github.com/pchlenski/misdeed.

---

The behavior of the microbiome, greatly elucidated by improvements in genome sequencing and data analysis, is generating considerable research interest. For instance, the Human Microbiome Project (Turbaugh *et al*., 2007) endeavors to collect data on a mass scale to investigate the role of the microbiome in the context of human health and disease. Despite improvements in sequencing, sample collection itself still incurs significant overhead and many niches remain understudied. Furthermore, microbial relative abundance data, the most typical form of data collected in such studies, has a number of properties that make classical statistical analysis challenging: it is longitudinal, compositional, noisy, and stochastic (Antoine *et al*., 2019). The genetic power calculator (Purcell *et al*., 2003) streamlined research in statistical genetics by facilitating closed-form power analysis of hypothetical studies. Similarly, several tools help design microbiome studies: Web-GLV (Kuntal *et al*., 2019) enables researchers to visualize the dynamics of microbial systems using assumed ecological parameters, and Mattiello *et al*. (2016) provide a power calculator for case-control studies on microbial ecosystems near equilibrium. The generalized Lotka-Volterra (gLV) modeling assumptions underlying such tools are often used to generate synthetic data when designing inference methods for microbial relative abundance data (Joseph *et al*., 2020). Here we present MiSDEED: the Microbial Synthetic Data Engine for Experimental Design, a flexible tool for generating synthetic longitudinal data from dynamic simulated ecosystems. These simulations can help investigators allocate resources in their study design, determine how to salvage underpowered studies, and design novel inference techniques with known ground truth.

MiSDEED’s synthetic data generator (drawn in Figure 1) samples reads from probability distributions governed by gLV dynamics over a discrete set of time points *T*. Each generator has a set *I* of nodes which may represent different data types (e.g. metagenomics and metabolomics measurements of the same system) or two interacting ecosystems with the same data type. Each node *i* ∈ *I* is initialized with a fixed dimensionality *d_i_*, a vector of growth rates 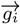, and an initial abundance vector 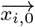. A generator also has up to |*I*|^2^ pairwise directed interactions between nodes. An interaction **A** between some nodes *i* and *i*′ is a matrix of dimension *d_i_* × *d_i′_*. Finally, the generator has a set *J* of interventions which may be applied to any node *i* ∈ *I* such that each intervention *j* has a vector 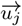 of intervention magnitudes and another vector 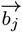 of responses to the intervention. If intervention *j* is applied to node *i*, then 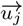 should have *T* dimensions and 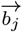 should have *d_i_* dimensions. To aid in parameter selection, MisDEED also contains convenience functions to initialize random but stable interaction matrices (Allesina and Tang, 2012) or to infer gLV parameters from known absolute abundance data (e.g. from a small pilot dataset) (Stein *et al*., 2013). Once generator parameters have been set, synthetic data can be produced in one of three ways: as a single timecourse, as multiple timecourses from varying initial conditions, or as multiple timecourses following a case-control split. In each case, the generator numerically solves the following equations with a biological noise term 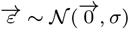

**Fig. 1.**
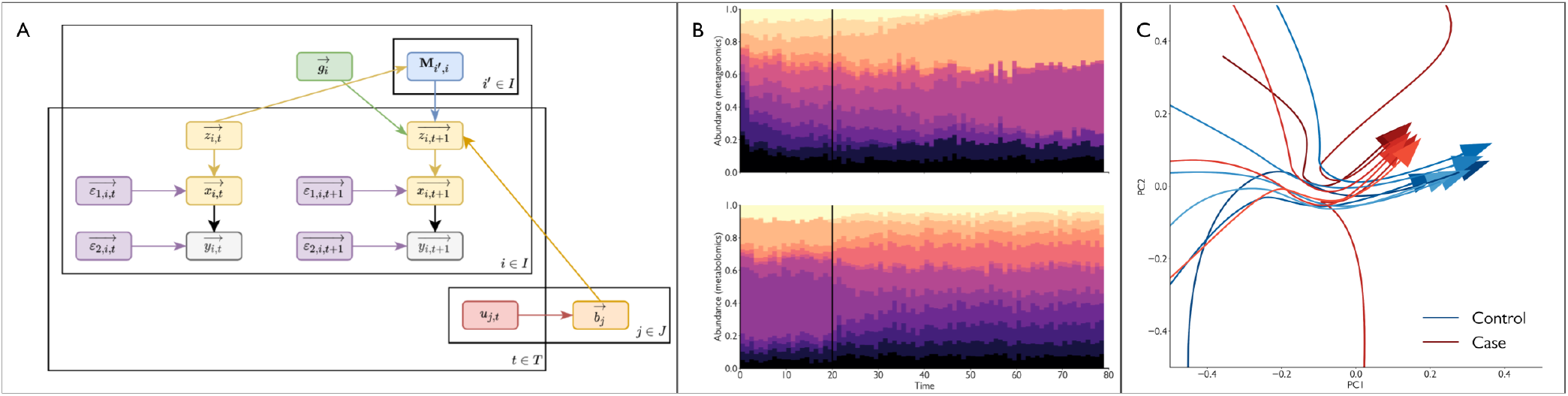
(A) The graphical model underlying MiSDEED’s synthetic data engine. The 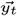 vectors are sampled from a multinomial distribution parameterized by the total number of reads and the probability vector 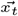. (B) Simulated metagenomic (top) and metabolomic (bottom) relative-abundance timecourses with an intervention at *t* = 20 (black line). This intervention affects metabolite abundances directly and propagates into the metagenomics node gradually via metabolomics-metagenomics interactions. (C) 12 noiseless PCA-transformed case (red) and control (blue) metagenomic trajectories show how interventions induce convergence to distinct fixed points.

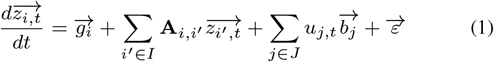

Each timecourse contains three derived matrices of synthetic data: *Z* (latent absolute abundances), *X* (latent relative abundances/probabilities), and *Y* (relative abundances sampled from *Y*).

MiSDEED is designed to be used as a standalone command line tool. The MiSDEED repository also contains the Python package underlying MiSDEED, a handful of utility scripts to support data visualization and learning gLV parameters, and a set of Jupyter notebooks showing common uses of the MiSDEED Python package. MisDEED can produce, save, and plot large amounts of synthetic data with varying initial conditions and model assumptions. To support power analysis, many variables can freely be changed by the user. These are listed in Table 1.

**Table 1.** Variable parameters in MiSDEED.

| Category | Parameters |
| --- | --- |
| Generator | Number of time points, number of nodes, node names, node dimensions, time to first sample |
| Random interaction matrices | $C$ (connectivity), $d$ (negative self-interaction size) $\sigma$ (multivariate normal variance), $\rho$ (multivariate normal correlation) |
| Custom gLV parameters | Interaction matrices, growth rates, initial abundances, interventions and intervention responses |
| Synthetic data generation | Biological noise variance, number of reads, time step size, downsampling rate |
| Multiple samples | Number of individuals, probability of 0-valued initial abundances |
| Case-control | Case-control ratio, intervention node, intervention effect size |

As an example use case, one may use a community matrix and growth rates learned from a pilot dataset and initialize ‘metagenomics’ and ‘metabolomics’ nodes such that the latter has no intrinsic growth rates, weak self-interactions, but strong interactions with the ‘metagenomics’ node according to some a priori assumptions. Perturbing metabolite abundances directly, the user may investigate how many patients must be enrolled in order to distinguish reliably between samples with and without this perturbation applied.

MiSDEED is a flexible framework for rapidly generating large amounts of realistic microbial trajectory data, thereby facilitating study design without assuming the system being studied is at equilibrium. Future development will focus on expanding code-free interfaces to MiSDEED; more flexible modeling assumptions for broader use cases, including nonuniformtime points, individual variation in interaction matrices and growth rates, and population clusters; alternatives to gLV-based modeling for dynamics like mutualism; and investigation into the value of MiSDEEDgenerated data for transfer learning and algorithm development.

## Acknowledgments

The work was supported by NIH/NCI Grant No. U54CA209997 Driving Biological Projects and Columbia University’s 2020/2021 Data Science Institute Seed Grant.

